# Pathogenic variants in Crx have distinct cis-regulatory effects on enhancers and silencers in photoreceptors

**DOI:** 10.1101/2023.05.27.542576

**Authors:** James L. Shepherdson, Ryan Z. Friedman, Yiqiao Zheng, Chi Sun, Inez Y. Oh, David M. Granas, Barak A. Cohen, Shiming Chen, Michael A. White

## Abstract

Dozens of variants in the photoreceptor-specific transcription factor (TF) CRX are linked with human blinding diseases that vary in their severity and age of onset. It is unclear how different variants in this single TF alter its function in ways that lead to a range of phenotypes. We examined the effects of human disease-causing variants on CRX *cis*-regulatory function by deploying massively parallel reporter assays (MPRAs) in live mouse retinas carrying knock-ins of two variants, one in the DNA binding domain (p.R90W) and the other in the transcriptional effector domain (p.E168d2). The degree of reporter gene dysregulation caused by the variants corresponds with their phenotypic severity. The two variants affect similar sets of enhancers, while p.E168d2 has stronger effects on silencers. *Cis*-regulatory elements (CREs) near cone photoreceptor genes are enriched for silencers that are de-repressed in the presence of p.E168d2. Chromatin environments of CRX-bound loci were partially predictive of episomal MPRA activity, and silencers were notably enriched among distal elements whose accessibility increases later in retinal development. We identified a set of potentially pleiotropic regulatory elements that convert from silencers to enhancers in retinas that lack a functional CRX effector domain. Our findings show that phenotypically distinct variants in different domains of CRX have partially overlapping effects on its *cis*-regulatory function, leading to misregulation of similar sets of enhancers, while having a qualitatively different impact on silencers.

## Introduction

The transcription factor (TF) Cone-Rod Homeobox (CRX) plays a key role in the differentiation and maintenance of photoreceptors (Chen et al. 1997; Furukawa et al. 1997; Freund et al. 1997; Swaroop et al. 2010). Variants in *CRX* have been shown to cause several inherited retinal diseases, including Retinitis Pigmentosa (RP), Cone-Rod Dystrophy (CoRD), and Leber Congenital Amaurosis (LCA) (Evans et al. 1994; Swain et al. 1997; Sohocki et al. 1998; Swaroop et al. 1999; Rivolta et al. 2001; Perrault et al. 2003; Ziviello et al. 2005; Nichols et al. 2010; Huang et al. 2012; Koyanagi et al. 2019; Fujinami-Yokokawa et al. 2020; Ng et al. 2020). *CRX* is the only gene implicated in all three of these diseases (Berger et al. 2010; Tran and Chen 2014). Retinopathies associated with *CRX* present with different phenotypes, varying in their cone vs. rod predominance, severity, and age of onset. Some *CRX* variants have been shown to cause severe dominant disease, while others are pathogenic only in a recessive context (Rivolta et al. 2001; Huang et al. 2012; Hull et al. 2014; Fujinami-Yokokawa et al. 2020; Nishiguchi et al. 2020; Kim et al. 2023).

The variable phenotypes of different pathogenic variants in *CRX* are likely due to differences in their effects on gene expression (Chen et al. 2002; Tran and Chen 2014). Studies of mouse models carrying retinopathy-associated variants p.R90W or p.E168d2 show that major changes in gene expression are evident by P10, during the critical period of photoreceptor differentiation (Swaroop et al. 2010) and before retinal degeneration can be observed (Tran et al. 2014; Ruzycki et al. 2015; Zheng et al. 2023). These studies also show that the two variants have graded effects on gene expression that correspond with the severity of their retinal phenotypes. Genes affected by p.R90W are also misregulated by the more phenotypically severe p.E168d2, which affects additional genes. However, changes in gene expression are an indirect measure of variant effects on CRX function since RNA levels can also be altered by other means, such as changes in concentrations of other TFs or changes in transcript stability. Pathogenic CRX variants could alter gene expression by affecting DNA binding or protein-protein interactions at thousands of different *cis*-regulatory elements (CREs), depending on the number, arrangement, and affinity of TF binding sites at each CRE. CRX is known to interact with other photoreceptor TFs and transcriptional co-factors (Mitton et al. 2000; Yanagi et al. 2000; Srinivas et al. 2006; Roduit et al. 2009; Peng et al. 2005; Hlawatsch et al. 2013; Sun and Chen 2023; Peng et al. 2007; Langouët et al. 2022). However, changes in *cis*-regulatory activity caused by pathogenic variants are not easily predictable from detailed biochemical knowledge of specific protein-DNA and protein-protein interactions. To understand how pathogenic variants in CRX affect its function as a TF, it is important to consider how these variants alter *cis*-regulatory activity across a broad range of CRX-bound CREs.

Reporter gene assays offer a more direct measure of CRX function at its target CREs. We previously used Massively Parallel Reporter Assays (MPRAs) in wild-type and *Crx^-/-^* mouse retinal explants to measure the activity of libraries of genomic and synthetic DNA sequences that are bound by CRX *in vivo* or contain CRX-binding motifs (White et al. 2013; White et al. 2016; Friedman et al. 2021). These studies revealed important TF motifs and other DNA sequence features that distinguish different classes of CRX-bound CREs, such as enhancers and silencers. Here we sought to understand how two biochemically and phenotypically distinct CRX variants alter its *cis*-regulatory function by determining how the variants affect the activities of different classes of CRX-bound CREs.

## Results

### Biochemically distinct CRX variants differ in the severity of their effects on CREs

We tested an MPRA library of CRX-bound CREs in retinal explants from mice that were either wild-type for *Crx* or carrying prototypical pathogenic *Crx* variants: the p.R90W single residue substitution variant and the p.E168d2 frameshift variant (Fig. 1A). Arginine 90 lies in the DNA-binding homeodomain of CRX, and the R90W variant has been shown to severely weaken binding to target CREs and reported to cause mild late-onset CoRD or LCA depending on zygosity (Swaroop et al. 1999; Hull et al. 2014; Tran et al. 2014; Fujinami-Yokokawa et al. 2020; Zheng et al. 2023). Glutamic acid 168 is located within the WSP domain of the transcriptional effector region, and frameshift into the +2 reading frame is predicted to lead to a stop codon within 3 residues. Because this stop codon is located within the final exon, the *p.E168d2* transcript is not predicted to undergo nonsense-mediated decay (Karousis and Mühlemann 2019). Although much of the transcriptional effector domain is lost, the p.E168d2 variant retains the ability to bind DNA (Chau et al. 2000; Tran et al. 2014). The p.E168d2 variant has been associated with dominant LCA in patients (Jacobson et al. 1998, Freund et al. 1998, Tran et al. 2014). Mice carrying these variants exhibit significant changes in gene expression by P10, and these changes result in developmental defects and eventual photoreceptor degeneration as early as three to four weeks of age (Tran et al. 2014; Ruzycki et al. 2015).

**Figure 1:**
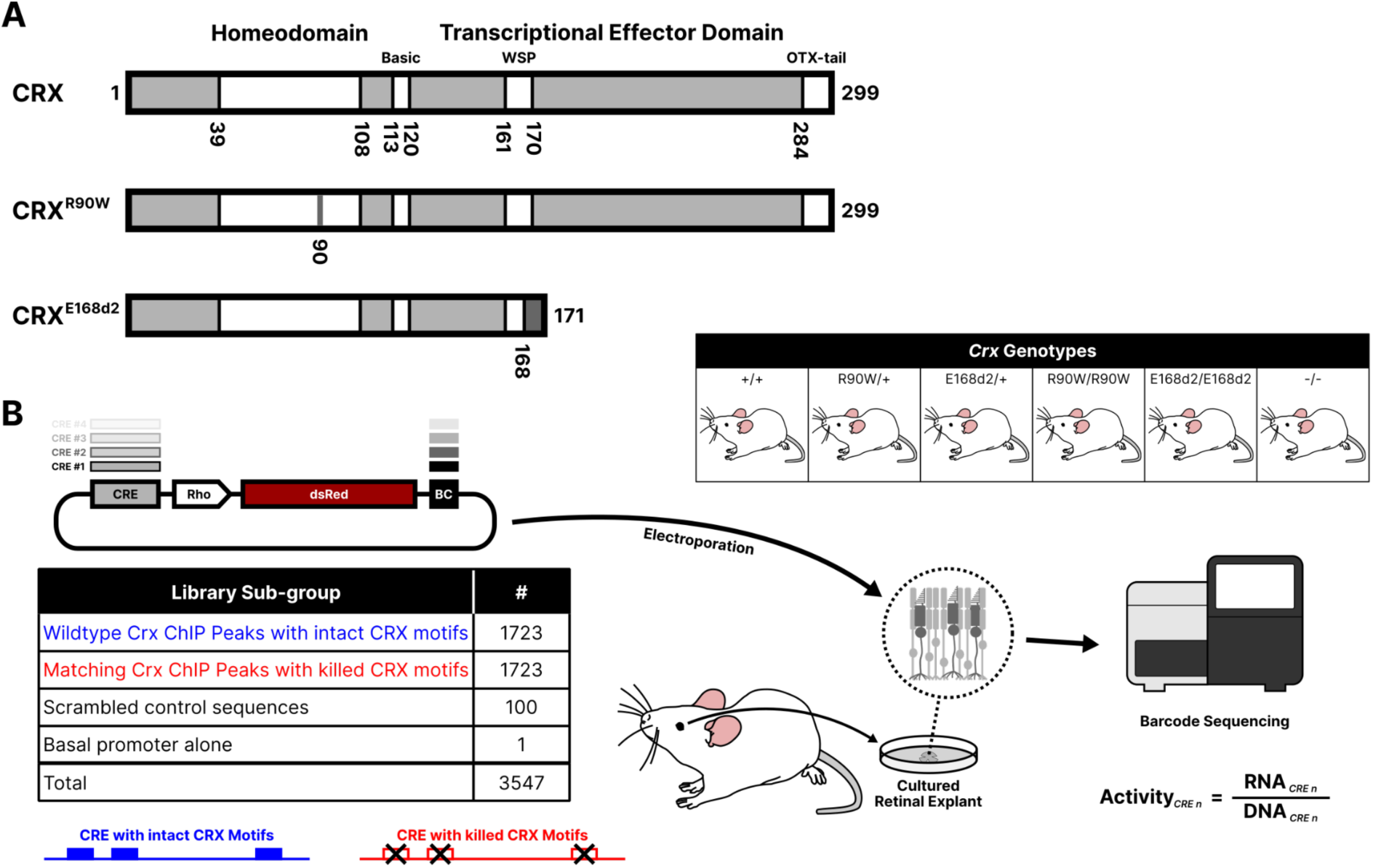
(**A**) Schematics of wild-type CRX protein and the p.R90W and p.E168d2 pathogenic variants, containing a point DNA binding domain mutant or resulting in truncation of the transcriptional effector domain, respectively. (**B**) Outline of the experimental procedure for constructing and testing the CRE libraries. A library of CRE sequences is cloned upstream of a rhodopsin (“Rho”) driving expression of the dsRed fluorescent protein, with each CRE marked by a unique sequence barcode (“BC”) in the 3’ UTR. Each library was electroporated into retinal explants with the six indicated *Crx* genotypes. RNA was collected from the retinas, and transcript counts for each element were measured and normalized to abundance in the input DNA library to calculate transcriptional activity.

To construct an MPRA library that tests the effects of variants on the *cis*-regulatory function of CRX, we selected 1,723 genomic CRE sequences that are bound by CRX as measured by ChIP-seq in whole adult mouse retina (Corbo et al. 2010; Ruzycki et al. 2018). CREs were selected to sample CRX-bound sites from three different chromatin environments that were previously measured at P14 (Ruzycki et al. 2018). For each CRE, we included a matching CRX motif mutant, in which all copies of the CRX motif were abolished using a mutation known to abrogate TF binding (White et al. 2013; Lee et al. 2010) (Fig. 1B). We cloned these 3,446 elements in a library with each CRE located upstream of a minimal promoter derived from the endogenous *rhodopsin* (*Rho*) locus driving expression of the fluorescent protein dsRed (Kwasnieski et al. 2012). Each CRE could be uniquely identified by a short sequence barcode located in the 3’ untranslated region of the dsRed transcript. We also included 100 scrambled elements as negative controls, with sequence composition matching that of the average CRE but with no detectable CRX binding motifs as called by FIMO using a p-value cutoff of 0.0025 (Grant et al. 2011).

We introduced this library via electroporation into P0 retinal explants from wild-type *C57BL/6J* mice (*+/+*), mice heterozygous or homozygous for the *p.R90W* (*R90W/+* and *R90W/W*) or *p.E168d2* (*E168d2/+* and *E168d2/d2*) variant, and *Crx* knockout mice (-/-) (all on *C57BL/6J* background) (Fig. 1B). After extracting RNA at P8, we sequenced CRE barcodes and normalized read counts to the abundance of each element in the input plasmid library (retina RNA read count / plasmid DNA read count) to compute each element’s transcriptional activity. R^2^ values between biological replicates were high, ranging from 0.90 in the +/+ background to ∼0.75 in the homozygous mutant backgrounds (Supplemental Fig. S1). We classified each CRE into one of five functional classes based on their transcriptional activity relative to the basal promoter alone: strong enhancer (significant increase in transcriptional activity relative to basal and greater than the 95th percentile of the scrambled sequences), weak enhancer (significant increase in transcriptional activity relative to basal and less than or equal to the 95th percentile of the scrambled sequences), inactive (no significant transcriptional activity difference compared to basal), weak silencer, and strong silencer (using the same criteria and cutoffs as for enhancers, but with significantly less transcriptional activity than basal) (Friedman et al. 2021).

We first evaluated the global *cis*-regulatory effects of the variants by comparing CRE expression from each variant mouse line to retinas from *+/+* mice (Fig. 2A and B). We observed an increasingly pronounced global dysregulation of CRE activity that correlated with increasing phenotypic severity. Phenotypic severity was previously shown to correspond with the identity and zygosity of the mutants (Tran et al. 2014). Little retinal impairment is observed in *R90W/+* mice, while progressive rod dystrophy occurs in *E168d2/+* mice. Unlike the heterozygotes, all homozygous mutant mice are blind, with *E168d2/d2* mice exhibiting the most severe photoreceptor degeneration among the homozygous mutant mice. Consistent with the relative phenotypic severities of the genotypes, we found that CRE activity in retinas from mice carrying the least-severe *R90W/+* genotype was tightly correlated with activity in corresponding *+/+* retinas (R^2^ = 0.85). In contrast, CRE activity in retinas with the severe *E168d2/d2* genotype correlated poorly with matched *+/+* measurements (R^2^ = 0.25).

**Figure 2:**
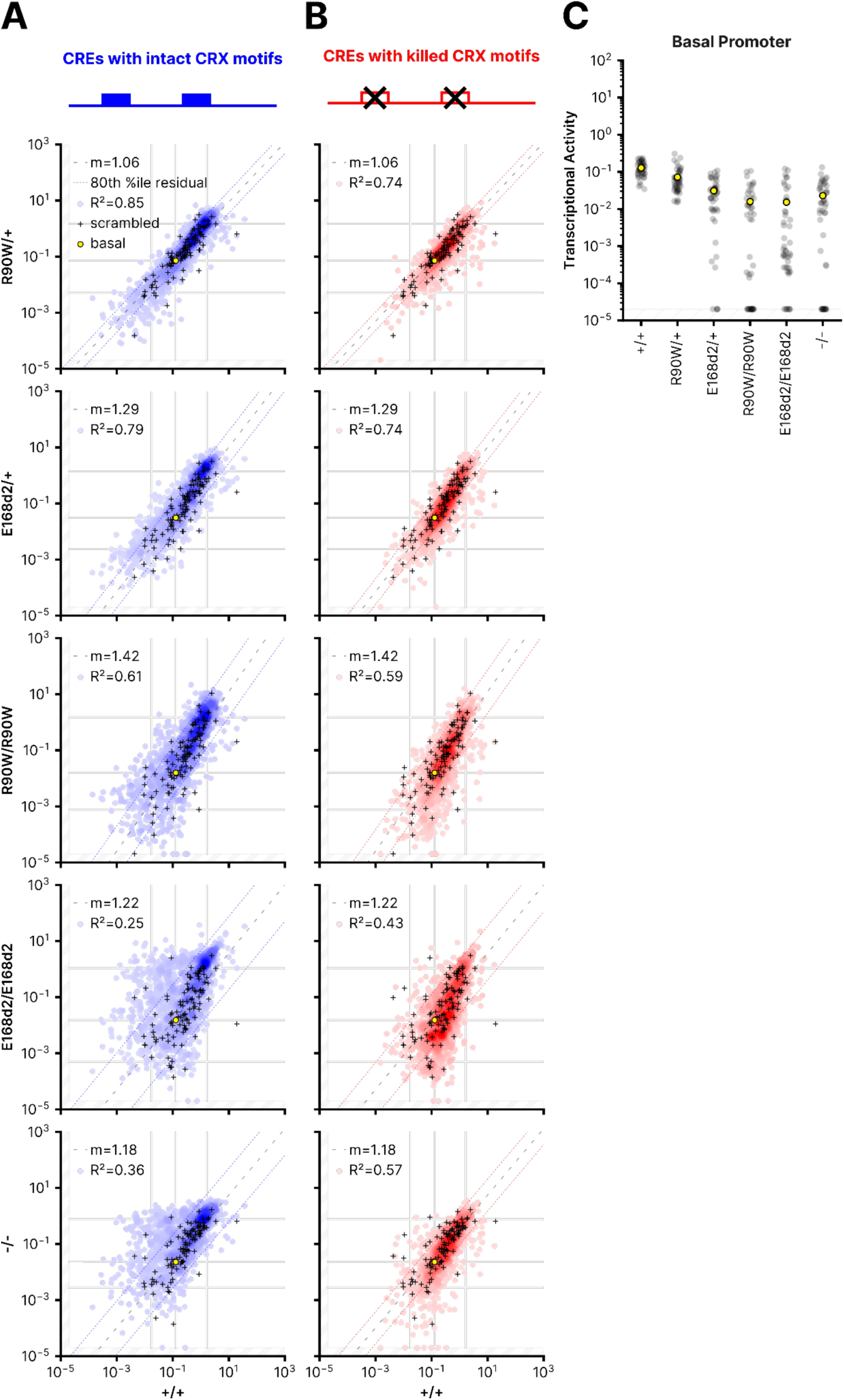
(**A,B**) Correlations of the transcriptional activity of each CRE with intact CRX motifs (**A**) or CRE with killed CRX motifs (**B**) in the indicated genotypes. Superimposed black cross symbols indicate the transcriptional activity of the 100 scrambled control sequences, and the yellow circle shows the activity of the basal rhodopsin promoter alone. Using the scrambled control sequence to establish an empirical null distribution of CRE activity, we performed robust regression using a trimmed mean M-estimator (fit plotted as a dashed line) for each genotype comparison. The outer dotted lines indicate the 80th percentile of the residuals of the library CREs relative to the regression of the scrambled sequences. The outer vertical and horizontal lines indicate the thresholds for the “strong silencer” and “strong enhancer” classes, while the inner horizontal and vertical lines correspond to the basal rhodopsin promoter alone. (**C**) Activity of the basal promoter construct in each of the indicated genotypes. Each black dot indicates a different unique sequence barcode; the mean across barcodes is shown as a yellow dot. In all panels, transcriptional activity was adjusted for visualization purposes using a fixed pseudocount of 2e^-5^; the out-of-bound region is indicated by the light gray stripes.

The *Rho* minimal promoter contains CRX binding sites and thus its intrinsic transcriptional activity is also altered by CRX variants. We quantified this effect by calculating the regression slope for each mutant genotype compared with wild-type, using a trimmed mean M-estimator based on the 100 scrambled control sequences (Fig. 2A, M-estimator fit is plotted by the dashed line). The slope increasingly deviates from 1 in the more severe genotypes, indicating that a decline in basal promoter activity contributes to the dysregulation of the reporter genes. The M-estimator values are consistent with an observed decrease in the activity of the *Rho* minimal promoter alone, as measured by MPRA barcodes that tag the basal promoter (Fig. 2C). Together, the *R^2^* and the M-estimator value show that the degree of global CRE dysregulation and the reduction in basal promoter levels both correspond with the phenotypic severity of the mutants. When CRX motifs in the CREs are abolished, the correlation between the *E168d2/d2* and *-/-* genotypes and +/+ retinas improves (Fig. 2B). This shows that much of the dysregulation in these genotypes is mediated by CRX motifs. Across all genotypes, most CREs alter their activity when CRX motifs are abolished (Supplemental Fig. S2). These results indicate that the dysregulation of the reporter gene library is a direct consequence of altered CRX function.

We examined specific classes of CREs and observed qualitatively different responses among strong and weak enhancers to the mutant genotypes. In *R90W/+* retinas, 134 (13%) strong and weak enhancers lost substantial activity, defined as a reduction to a lower activity class (Supplemental Fig. S3A). More enhancers lost activity in the more severe *R90W/R90W* (n = 164, 16%) and *E168d2/d2* genotypes (n = 212, 21%). A subset of weak enhancers exhibited amplified transcriptional activity relative to the basal promoter in the mutant genotypes. 114 weak enhancers (13%) increased activity relative to the *Rho* minimal promoter in *R90W/+* retinas, while 150 (18%) and 195 (22.9%) were amplified in *R90W/R90W* and *E168d2/d2* retinas, respectively (Supplemental Fig. S3A). This increase in activity relative to basal levels was not due to an absolute increase in transcriptional activity, but was instead due to a decrease in basal activity in the more severely affected genotypes (Fig. 2A and C). Thus, these CREs retained some independent transcriptional activity as the basal promoter activity decreased in the mutant genotypes. The persistent activities of these CREs suggests that they are only weakly affected by a loss of CRX function and are able to drive robust activation from a weakened basal promoter. We searched for TF binding motifs enriched in subsets of enhancers that responded differently to the mutant genotypes, but found no motifs that were strongly enriched in any subset of enhancers (Methods). This suggests that the responses of enhancers to CRX variants are not defined by the presence or absence of particular motifs for cooperating TFs, but instead by a more complex *cis*-regulatory grammar defined by multiple TFs that interact with CRX.

The most striking difference between the p.R90W and p.E168d2 variants was in their effect on silencers in the homozygous mutant genotypes. Some silencers were moderately de-repressed in *R90W/W* retinas, but a more extensive loss of repression among silencers occurred in *E168d2/d2* and *-/-* retinas (Fig. 2A). This suggests that the transcriptional effector domain of CRX, absent in *E168d2/d2* and *-/-* retinas, is critical for CRX function at silencers. The extensive de-repression observed in *E168d2/d2* retinas relative to *R90W/W* retinas is consistent with results from a prior RNA-seq analysis of these mouse models (Ruzycki et al. 2015). In this study, more gene upregulation occurred in *E168d2/d2* retinas (248 of 673 differentially expressed genes, 37%) compared to *R90W/R90W* retinas (70 of 265 differentially expressed genes, 26%, p = 0.003, Fisher’s exact test). By RNA-seq analysis alone, upregulation of genes in the *E168d2/d2* genotype could be an indirect effect of loss of CRX function, possibly due to changes in the levels of other TFs. However, our finding that silencers are de-repressed in this genotype suggests that gene upregulation is a direct effect caused by the loss of CRX-mediated silencing.

### CREs near cone photoreceptor genes are enriched for silencers

A previous RNA-seq analysis found that genes enriched in rod photoreceptors were largely downregulated in mouse retinas carrying the p.R90W and p.E168d2 variants, while genes enriched in cone photoreceptors were frequently upregulated (Ruzycki et al. 2015). Because rod photoreceptors comprise 80% of the cells in mouse retina, gene expression analyses of whole retina primarily capture gene expression changes in rod photoreceptors, and thus the loss of expression of rod genes and the derepression of cone genes in the mutant genotypes reflect changes in CRX function in rod photoreceptors.

We examined whether changes in CRE activity in the mutant genotypes recapitulated changes in rod and cone gene expression. While the MPRA library was designed to sample CREs from different chromatin environments, and not by proximity to known photoreceptor genes, we identified 183 CREs in the library within 100 kbp of a transcription start site (TSS) of 241 previously identified rod- or cone-enriched genes. 117 CREs in the MPRA library were near rod genes, while 66 were near cone genes. CREs near cone genes were highly enriched for silencers compared with CREs near rod genes (Fig. 3A, cone CREs: n = 25 (37.8%); rods CREs: n = 16 (13.7%)), while rod genes were enriched for enhancers (rod CREs: n = 76 (65.0%), cone CREs: n = 37 (56.1%), p = 0.022, chi-square test for independence). Correspondingly, CREs near cone genes were more likely to be de-repressed in the mutant genotypes, particularly in *E168d2/d2* and *-/-* that lack the CRX effector domain (Fig. 3B and C). While changes in the expression of a gene are the cumulative result of changes in the activity of multiple CREs, these results show that an episomal assay of CRE activity partially recapitulates changes in downstream gene expression. One role of CRX is to direct repression of cone genes in rod photoreceptors in cooperation with other TFs such as Nr2e3 (Peng et al. 2005). Our results suggest that this repressive role requires the CRX effector domain, and that the loss of repressive activity of the p.E168d2 variant may explain its more severe phenotype.

**Figure 3:**
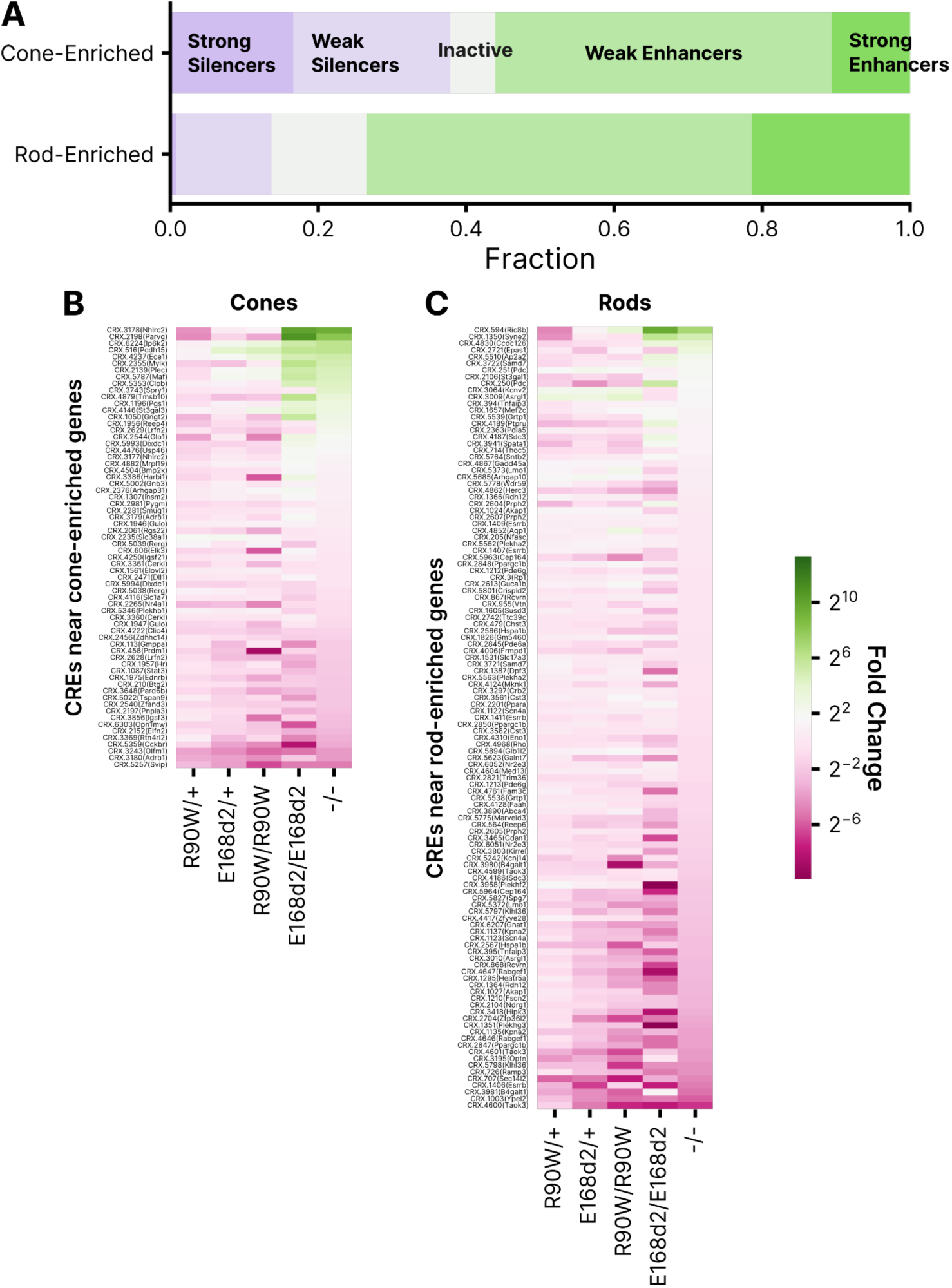
(**A**) Distribution of activity classes in +/+ retina for 66 and 117 CREs near cone- or rod-enriched genes, respectively (Ruzycki et al. 2015). (**B**) Fold change relative to +/+ of 66 CREs near cone-enriched genes. (**C**) Fold change relative to +/+ of 117 CREs near rod-enriched genes. In (B) and (C), rows are sorted by fold change in -/-.

### A subset of CREs convert from silencers to enhancers in genotypes lacking the CRX effector domain

Most CREs exhibit a graded response across the CRX mutants, with the effects differing primarily in their severity (Fig. 4A). However, a subset of silencers exhibit not only loss of repressive activity, but a gain of enhancer activity in *E168d2/d2* and *-/-* retinas (Fig. 4A, upper arrow). We identified 234 sequences classified as weak or strong silencers in wild-type retinas that become weak or strong enhancers in the *E168d2/d2* and *-/-* retinas, which lack the CRX transcriptional effector domain (Fig. 4B, outlined by the black square, Supplemental Fig. S3 A and C). Comparing CREs that convert from silencers to enhancers and their matched motif mutants across all genotypes, we observed an inversion of the effect of the CRX motif mutations in the *E168d2/d2* and *-/-* genotypes (Fig. 4C, Supplemental Fig. S3B). Specifically, mutating the CRX motifs caused de-repression in genotypes expressing CRX with an intact transcriptional effector domain (+/+, *R90W/+*, *E168d2/+*, *R90W/W*), while the same motif mutations caused loss of activation in genotypes that lack the CRX transcriptional effector domain (*E168d2/d2*, *-/-*). This effect does not occur in silencers that are merely de-repressed in the *E168d2/d2* and *-/-* genotypes (Fig. 4C). This demonstrates that these particular CRX motifs are pleiotropic, mediating repression in the presence of wild-type CRX (when these CREs act as silencers) and mediating activation when the CRX transcriptional effector domain is lost in the *E168d2/d2* and *-/-* genotypes (when these CREs act as enhancers). Because this effect occurs in *Crx^-/-^* retinas, these pleiotropic sites must be bound by another TF in the absence of CRX. That TF is possibly the CRX ortholog OTX2 (Koike et al. 2007).

**Figure 4:**
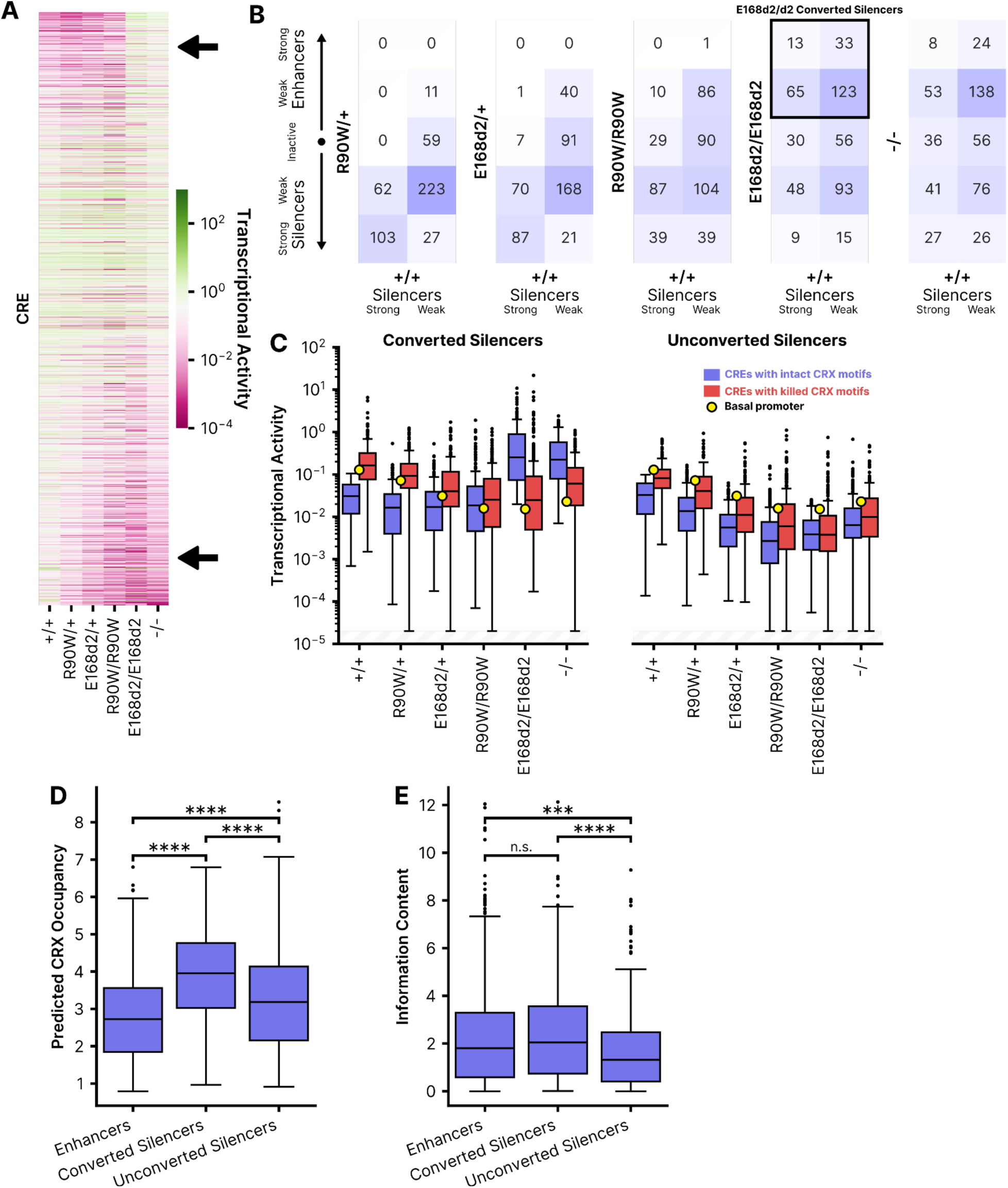
(**A**) Heatmap comparing the transcriptional activity of each CRE across the six CRX genotypes. In the top right (upper arrow), note a group of CREs with generally low activity in all but the E168d2/d2 and -/- genotypes. Along the bottom (lower arrow) note the stepwise reduction in activity with increasing phenotypic severity. CREs are sorted by the ratio of transcriptional activity in +/+ and -/- genotypes. (**B**) Classifications across genotypes for CREs classified as strong silencers (left column of each heatmap) or weak silencers (right column) in wildtype retinas. (**C**) Transcriptional activity of wildtype or motif mutant CREs across genotypes for 234 silencers in +/+ that convert to enhancers in E168d2/d2 (“Converted”), or the remaining 251 silencers in +/+ that do not convert to enhancers in E168d2/d2 (“Unconverted”). The yellow dot indicates the transcriptional activity of the basal promoter alone in each genotype. (**D**) Predicted CRX occupancy of the same silencer subgroups as in panel C and CREs that act as enhancers in +/+. (**E**) Information content of the same subgroups as panel D. *: p < 0.05, **: p < 0.01, ****: p < 0.0001 via two-sided Mann-Whitney-Wilcoxon test. In panel C, transcriptional activity was adjusted for visualization purposes using a fixed pseudocount of 2e-5; the out-of-bound region is indicated by the light gray stripes.

To further characterize these CREs, we examined two sequence features that we previously showed are able to distinguish CRX-bound enhancers from CRX-bound silencers (Friedman et al. 2021). Compared to enhancers, CRX-bound silencers tend to contain more copies of the CRX motif, which we quantify using a threshold-free motif scan called “CRX predicted occupancy” (White et al. 2013; White et al. 2016; Friedman et al. 2021). Enhancers are characterized by the presence of a more diverse collection of motifs for eight other lineage-specific TFs. We quantify the diversity of motifs in a CRE with an “information content” score, based on Boltzmann entropy, that considers the number and diversity of motifs in a CRE. Information content considers motifs or motif families from CRX/OTX2, NRL (leucine zipper), NEUROD1 (E-box binding factor), RORB (nuclear receptor), MAZ/SP4 (C2H2 zinc finger), MEF2D (MADS-box), RAX (Q50 homeodomain) and GFI1 (C2H2 zinc finger repressor). We previously reported that CRX-bound enhancers have higher information content than silencers (Friedman et al. 2021). In sum, our previous work shows that enhancers are characterized by high information content and modest CRX predicted occupancy, while silencers are characterized by low information content and high CRX predicted occupancy.

We determined the predicted CRX occupancy and information content of CREs that convert from silencers to enhancers in *E168d2/d2* and *-/-* retinas, and found that these CREs have features of both silencers and enhancers. Most silencers that convert to enhancers have higher predicted CRX occupancy than wild-type enhancers (Fig. 4D), which is consistent with our prior finding that silencers in general have a higher predicted CRX occupancy (Friedman et al. 2021). However, the CRX predicted occupancy of most silencers that convert to enhancers is even higher than the elevated predicted occupancy of the other silencers (median 3.95 vs. 3.19). Yet unlike other silencers, most silencers that convert to enhancers also have the higher information content that is characteristic of enhancers (Figure 4E). The information content of this group is roughly as high as the information content of +/+ enhancers (median 2.05 vs. 1.80). The mixed sequence characteristics of these CREs may explain why they convert from silencers to enhancers in *E168d2/d2* and *-/-* retinas. They function as silencers in wild-type retina due to their high CRX occupancy. Upon loss of the CRX transcriptional effector domain, they cease to function as silencers, possibly due to the loss of interactions with a CRX-interacting repressor complex. However, their high information content, reflecting the presence of motifs for other TFs, enables these CREs to function as enhancers in *E168d2/d2* and *-/-* retinas, in contrast to low-information content silencers that simply lose their repressive activity. The additional TFs recruited by the diverse binding motifs in these CREs may interact with OTX2 or some other TF that binds the pleiotropic CRX motifs in the *E168d2/d2* and *-/-* genotypes.

We searched for additional enriched motifs in these potentially bifunctional CREs using Simple Enrichment Analysis (SEA) (Bailey and Grant 2021). We found that CREs that convert from silencers to enhancers are enriched in motifs for a variety of paired-type homeodomain TFs. CRX and its ortholog OTX2 are K50 paired-type homeodomains that contain a lysine at residue 50 of the DNA binding domain which determines the specificity of these TFs for a TAATCC consensus sequence (Furukawa et al 1997). While CRX and OTX2 play critical roles specifically in photoreceptors and bipolar cells, other non-K50 paired-type homeodomain TFs are active in multiple retinal cell types (Hennig et al. 2008; Zagozewski et al. 2014). Silencers that convert to enhancers are enriched in motifs for TFs active in several retinal cell types: LHX-family TFs (horizontal and Mueller glial cells, p = 2.02×10^-8^); VSX1 and VSX2 (cone bipolar cells and retinal ganglion cells, p = 1.23×10^-5^); ISL2 (retinal ganglion cells, p = 2.87×10^-6^); VAX1 and other NK-family TFs (role in ventral retina dorsalization, p = 1.78×10^-5^); and RAX (photoreceptors and bipolar cells, p = 2.13 × 10^-4^). These CREs are also strongly enriched in homeodomain dimer motifs. Compared to CREs that remain enhancers and silencers in wild-type and *E168d2/d2* retina, CREs that convert from silencers to enhancers exhibit a 1.5-fold enrichment of homeodomain motif dimers (71.2% of bifunctional CREs contain at least 1 dimer versus 47.4% among CREs that remain silencers or enhancers, p = 1.78×1^-10^). Paired-type homeodomain TFs bind as both homo- and hetero-dimers, and the presence of these dimers in potentially bifunctional CREs may enable these elements to play regulatory roles in multiple cell types (Wilson et al. 1993; Rister et al. 2015; Hughes et al. 2018).

Finally, we asked whether potentially bifunctional CREs are co-bound by the co-repressor BCOR, a Polycomb repressive complex 1 (PRC1) factor that co-binds with CRX and is linked with early-onset retinal degeneration (Langouët et al. 2022). Only a minority of all silencers in wild-type retina overlapped a BCOR binding site (11.8%). However, silencers that convert to enhancers in *E168d2/d2* and *-/-* retinas were even less likely to be bound at P0 by BCOR (4.7%, chi square test p = 7.2×10^-4^), suggesting that these potentially bifunctional elements are distinct from other silencers, and that their silencing may not depend on PRC1.

### Chromatin environments of CRX-bound sequences are partially predictive of episomal MPRA activity

CREs in the MPRA library were selected from different chromatin environments as measured in wild-type retina. All library CREs were selected from sites of accessible chromatin (measured by ATAC-seq at P14) (Ruzycki et al. 2018) that overlapped CRX ChIP-seq peaks (measured in adult retina) (Corbo et al. 2010), but these sites varied in other chromatin annotations as determined in a previous study (Ruzycki et al. 2018). These annotations include 1) the presence or absence of histone marks H3K4me3 and H3K27ac in P14 wild-type retina, and 2) the loss or retention of chromatin accessibility in P14 *Crx^-/-^* retina (*i.e.*, whether accessibility is CRX dependent or independent). Based on these annotations, CRX-bound accessible sites were previously classified into four groups (Table 1) (Ruzycki et al. 2018). Group A sequences are proximal regulatory elements that are positive for the promoter-associated H3K4me3 epigenetic mark (Vermeulen et al. 2010) and H3K27ac, a mark associated with active CREs (Creyghton et al. 2010). Group A sequences typically lie within 5 kb of the nearest TSS. Group C includes active distal CREsdefined by the presence of H3K27ac without H3K4me3. Group D includes distal sites that lack both H3K4me3 and H3K27ac, indicating that they are accessible but inactive at P14. However, these sites show evidence of increased activation by 8 weeks (Ruzycki et al. 2018). Group B sites, defined by the presence of H3K4me3 and the absence of H3K17ac, were not included in our MPRA library. Group B includes fewer than 3% of P14 ATAC-seq peaks and we thus chose to focus on Groups A, C, and D. Within each group, CRX-bound sites exhibit either CRX-dependent or independent chromatin accessibility. Group A independent sequences are open in multiple non-retinal cell types and are accessible in the retina throughout postnatal development, while all other classes are primarily accessible only in the retina and vary in the developmental timing of their accessibility (Table 1).

**Table 1:** Features of chromatin environments of MPRA library sequences.

| Group | Histone Marks |  | Typical Distance to TSS | CRX-dependent Accessibility | Timing and specificity of accessibility | # CREs in MPRA library |
| --- | --- | --- | --- | --- | --- | --- |
|  | H3K4me3 | H3K27ac |  |  |  |  |
| A | + | + | 0-5 kb | Dependent | Retina-specific, late accessibility | 249 |
|  |  |  |  | Independent | Open in other cell types, always accessible | 632 |
| C | - | + | 5-500 kb | Dependent | Retina-specific, late accessibility | 311 |
|  |  |  |  | Independent | Retina-specific, early accessibility | 366 |
| D | - | - | 5-500 kb | Dependent | Retina-specific, late accessibility | 76 |
|  |  |  |  | Independent | Retina-specific, early accessibility | 89 |

We found that CREs taken from different chromatin environments exhibited systematic differences in episomal MPRA activity in P8 wild-type retina. Group A CREs whose accessibility is independent of CRX contain the highest proportion of enhancers, the lowest proportion of silencers, and exhibit on average the highest MPRA activity (Fig. 5A and B). This contrasts with Group A CRX-dependent CREs, which include fewer enhancers and more silencers, and exhibit significantly lower average MPRA activity (Fig. 5A and B, mean log2 RNA/DNA = -1.083 vs. - 0.477 for dependent vs. independent Group A, p = 5.0 × 10^-4^, independent two-sample t-test). The difference in the proportion of enhancers and silencers in the subclasses of Group A corresponds with the developmental timing of chromatin accessibility of these subclasses. Group A CRX-independent CRE are fully accessible by P1 and are more likely to be MPRA enhancers at P8. Group CRX-dependent CREs first become accessible around P7 and reach peak accessibility later in development (Ruzycki et al. 2018; Wilken et al. 2015), and they drive lower transcriptional activity on average in the MPRA at P8.

**Figure 5:**
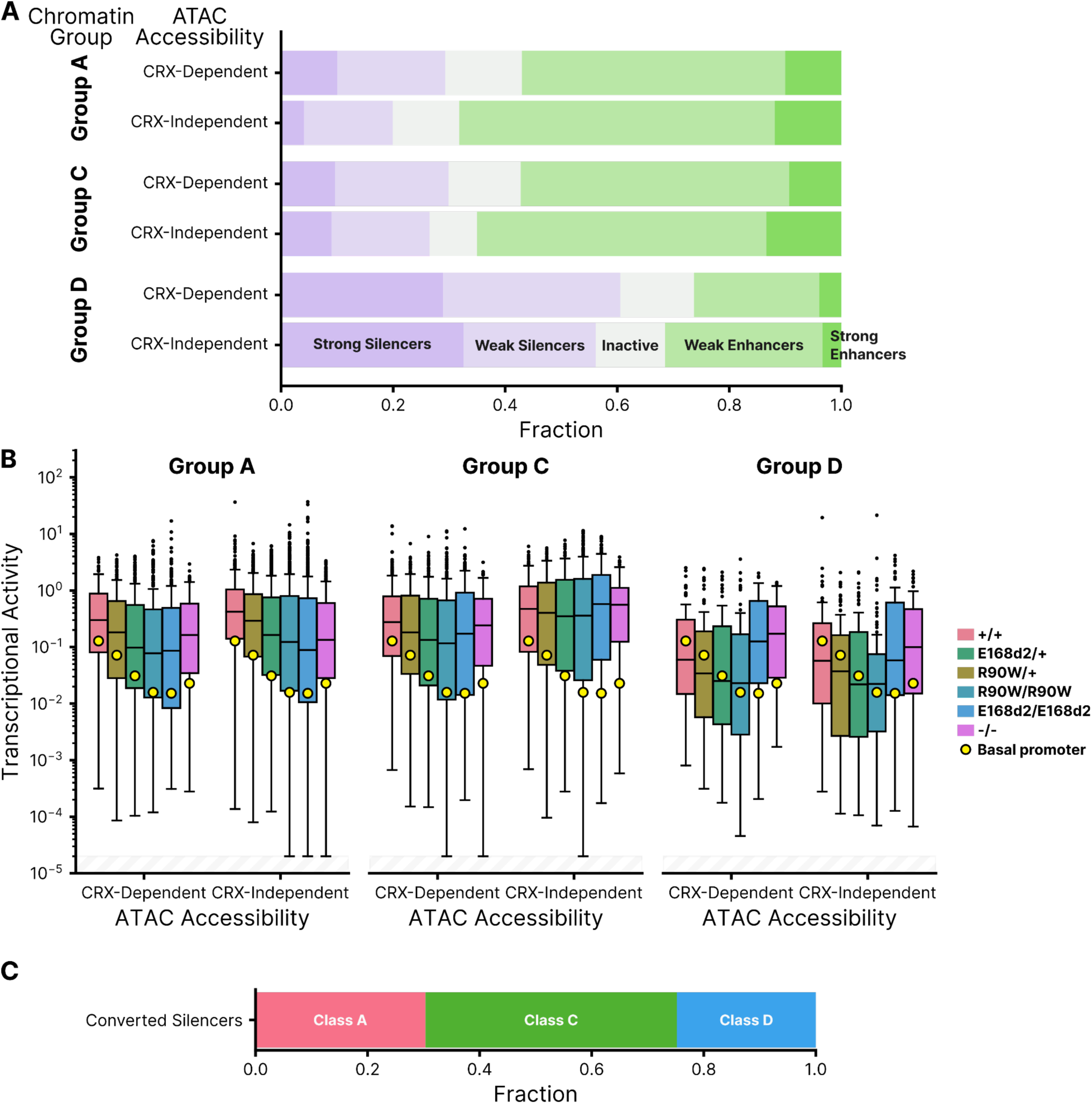
(**A**) Distribution of CRE activity classes in +/+ retina separated into groups based on chromatin annotations of the site of genomic origin and CRX-dependent ATAC accessibility (Ruzycki et al. 2018). Group A: H3K4me3+ H3K27Ac+, Group C: H3K4me3-H3K27Ac+, Group D: H3K4me3-H3K27Ac-. (**B**) Transcriptional activity of all CREs, separated by chromatin and ATAC group, across all genotypes. (**C**) Chromatin classification breakdown of E168d2/d2 converted silencers (elements classified as weak or strong silencers in +/+ that become weak or strong enhancers in E168d2/d2). In panel B, transcriptional activity was adjusted for visualization purposes using a fixed pseudocount of 2e^-5^; the out-of-bound region is indicated by the light gray stripes.

A similar trend was observed for the distal regulatory elements of Group C, which become accessible later in development. In Group C, CRX-independent CREs are partially accessible at P1 and almost fully accessible at P7, while CRX-dependent CREs only become partially accessible at P7 and reach full accessibility by 8 weeks. Consistent with these differences in the developmental timing of chromatin accessibility, Group C CRX-independent CREs have somewhat higher average MPRA activity than Group C CRX-dependent CREs (Fig. 5B, mean log2 RNA/DNA = -1.101 vs -0.672, p = 0.028, independent two-sample t-test), and they include a slightly higher proportion of strong enhancers (Fig. 5A). The majority of Group D CREs, which lack the H3K27ac mark of active regulatory elements, are silencers and inactive sequences and exhibit the lowest levels of MPRA activity of all chromatin environment groups (Fig. 5A and B). We also examined the distribution of converted silencers among the chromatin groups and found that these CREs are largely distal elements from Groups C and D (Fig. 5C, Supplemental Fig. S3D).

The correspondence between episomal MPRA activity and features of the genomic chromatin environment suggests that the sequence-encoded features responsible for MPRA activity also drive key chromatin changes at native genomic loci over the course of retinal development. Furthermore, the increased fraction of silencers among regulatory elements that become accessible later in development suggests that these later CREs may be repressive earlier in development, preventing premature expression of late developmental genes. Our results show that distal CREs which become active later in retinal development often behave as silencers when assayed at P8 by MPRA. These CREs are de-repressed in *E168d2/d2* and *-/-* retinas but not in *R90W/W* retinas (Fig. 5B, see especially Group D). This suggests that the more severe phenotype of *E168d2/d2* and *-/-* mouse models may be due in part to premature activation of distal CREs and a consequent disruption of developmentally timed gene expression.

### CRX mutants have no systematic effects on cooperativity between CRX binding sites

Included in the MPRA library are 100 CREs with exactly two CRX binding motifs. To test for effects of the p.R90W and p.E168d2 CRX mutants on homotypic cooperativity, we designed single and double CRX motif mutants for each of these CREs. If the two CRX motifs in a CRE act cooperatively, the sum of the effects of the single motif mutants will exceed the effect of the double motif mutant. To assess cooperativity we therefore computed the sum of the effects of the single motif mutants on MPRA activity, and divided that by the effect of the double motif mutant. A ratio greater than one indicates that the two motifs act cooperatively, while a ratio near one indicates no cooperativity. A ratio less than one could indicate either redundancy or anti-cooperativity between the two CRX motifs. We found that very few of the 100 tested CREs showed evidence of cooperativity between CRX sites, and none of the mutant *Crx* genotypes systematically increased or decreased cooperativity (Supplemental Fig. S4). While the p.R90W and p.E168d2 CRX variants may affect cooperative interactions between CRX and other TFs, or may affect CRX-CRX cooperativity in a very specific contexts, they have no systematic effects on homotypic cooperative interactions between CRX molecules.

## Discussion

We used MPRAs conducted in P0-8 whole retinal explants to understand how two biochemically distinct pathogenic variants alter the *cis*-regulatory function of CRX. In both humans and mouse models, pathogenic variants in CRX often lead to progressive photoreceptor degeneration beginning in young adulthood, but evidence from mouse models indicates that mis-regulated gene expression can be observed much earlier (Tran et al. 2014, Ruzycki et al. 2015; Zheng et al. 2023). In this study, we assayed CRX function when dysregulated *cis*-regulatory activity is already evident, but before photoreceptor degeneration has begun (Tran et al. 2014). All CREs included in the MPRA library were selected from genomic loci that are at least partially accessible at P14 and bound by CRX in adult retina (Ruzycki et al. 2018; Corbo et al. 2010), but they were sampled from a variety of chromatin environments and they vary in the developmental timing of their activation (Ruzycki et al. 2018; Wilken et al. 2015). Thus our MPRA study was designed to test CRX function across a broad range of its target CREs.

Consistent with a prior RNA-seq analysis of the same mouse models (Ruzycki et al. 2015), we observed graded degrees of global dysregulation in CRE activity that correspond with the severity of the retinal phenotypes of the mouse models (Figs. 2A and 4A). This suggests that the phenotypic differences between mice carrying p.R90W and p.E168d2 can be partially explained by the fact that p.E168d2 disrupts CRX *cis*-regulatory function at more CREs, particularly silencers. While p.R90W and p.E168d2 variants affect largely overlapping sets of wild-type enhancers (Fig. 4A, lower arrow), p.E168d2 causes extensive silencer de-repression that does not occur with p.R90W (Fig. 4A, upper arrow). Silencer de-repression also occurs in *Crx^-/-^* retina, showing that the repressive function of CRX at silencers requires the presence of its transcriptional effector domain, which has been classically considered an activation domain (Chen et al. 1997; Peng and Chen 2007). However, a recent study found that the transcriptional effector domain of human CRX is capable of both activation and repression, and that homeodomain TFs are enriched in such bifunctional effector domains (DelRosso et al. 2023). The ability of CRX to act as both a repressor and an activator is consistent with its role as a regulator of distinct gene expression programs in multiple photoreceptor cell types (Swaroop et al. 2010).

One key role of CRX is to direct repression of cone photoreceptor genes in rod photoreceptors (Peng et al. 2005), and a loss of repression of cone photoreceptor genes was observed by RNA-seq in whole *E168d2/d2* mouse retina (Ruzycki et al. 2015). Consistent with this, we found that CREs near cone photoreceptor genes were disproportionately silencers in our episomal MPRA, and these silencers were de-repressed in the *E168d2/d2* and *-/-* genotypes (Fig. 3). RNA-seq and MPRA analyses performed in whole retina largely reflect the *cis*-regulatory activity in rod photoreceptors, because ∼80% of the cells of mouse retinas are rods (Jeon et al. 1998) and rod make up 87% of plasmid-bearing cells in a whole retina MPRA (Kwasnieski et al. 2012; Zhao et al. 2023). Taken together, RNA-seq and MPRA results suggest that the loss of the CRX transcriptional effector domain leads to inappropriate activation of cone genes in rod photoreceptors early in development, and that this activation is a direct consequence of altered CRX function at its target silencers. Variants like p.E168d2 that truncate the CRX effector domain may abolish its ability to interact with other factors required for repression, such as Nr2e3 and Samd7 (Peng et al. 2005; Hlawatsch et al. 2013; Omori et al. 2017), and thereby impair rod differentiation.

Results of prior MPRA studies show that whether CRX acts as an activator or repressor at a particular CRE depends on local motif composition (White et al. 2013; White et al. 2016; Hughes et al. 2018; Friedman et al. 2021; Loell et al. 2023). We previously reported that CRX-bound silencers tend to have more copies of the CRX motif than enhancers do (White et al. 2016; Friedman et al. 2021), while enhancers contain more motifs for other photoreceptor TFs (Friedman et al., 2021). These features of the sequence composition of enhancers and silencers suggest a model that may explain why enhancers and silencers respond differently to the p.R90W and p.E168d2 variants (Fig. 6). At enhancers, which have moderate CRX occupancy, both variants cause a loss of active CRX, either by reducing DNA binding (p.R90W) or by loss of the effector domain (E168d2). At silencers, the presence of more copies of the CRX motif may cause higher CRX occupancy, which makes silencers less sensitive to the reduced DNA binding affinity of p.R90W. High CRX occupancy at silencers may be promoted by homotypic interactions mediated by the transcriptional effector domain, which is necessary for repression. We recently presented a neural network model of CRX-directed *cis*-regulation that supports a role for homotypic interactions (Loell et al. 2023). This model, trained on MPRA data from synthetic CREs with different combinations of photoreceptor TF binding sites, indicated that homotypic interactions between multiple copies of the CRX binding site were necessary to create repressive CREs. While the biophysical nature of these homotypic interactions is unknown, there is clear evidence that the presence of multiple copies of the CRX motif often leads to repression in both genomic and synthetic CREs (White et al. 2016; Friedman et al. 2021; Loell et al. 2023).

**Figure 6:**
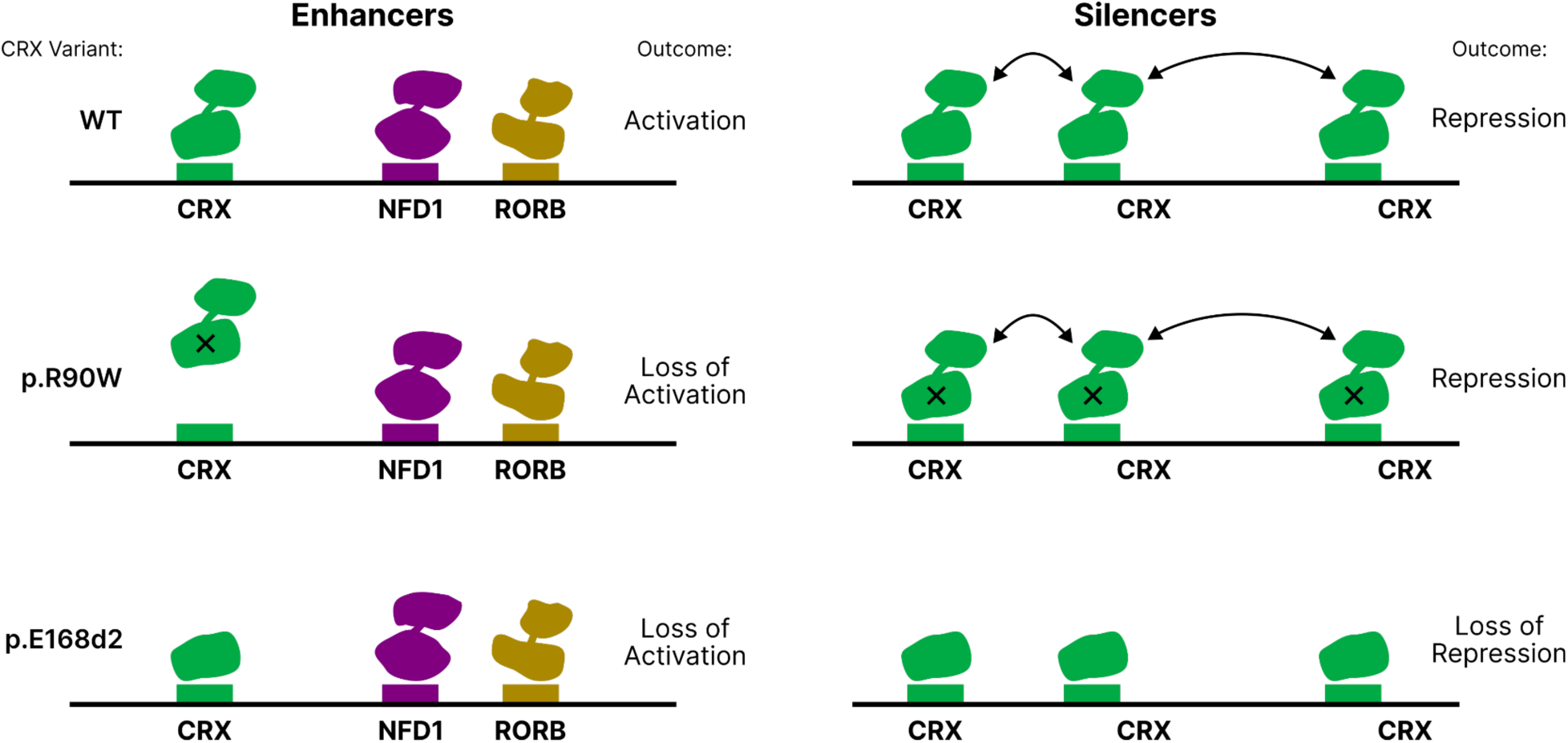
A model of differential enhancer and silencer sensitivity to *CRX* mutants. Silencers, with higher predicted CRX occupancy, engage in effector-domain mediated homotypic interactions that permit CRX to remain bound even when the DBD is weakened by the p.R90W variant. The p.E168d2 variant lacks the majority of the effector domain and cannot engage in these interactions. Enhancers have lower CRX predicted occupancy and engage in fewer homotypic interactions.

Characteristic sequence features of both silencers and enhancers are present in the potentially bifunctional CREs that we report here. These CREs, which are silencers in *+/+* retinas and convert to enhancers in *E168d2/d2* and *-/-* retinas, have especially high copy number of the CRX motif, as well as motifs for multiple other lineage-specific TFs (Fig. 4D and E). The ability of these silencers to convert to enhancers in genotypes that lack the CRX transcriptional effector domain may indicate that these CREs play a role as enhancers in other cell types or at other developmental time points in wild-type photoreceptors. The CRX binding sites in these CREs are clearly pleiotropic, being necessary for repression in *+/+* retinas and for activation in *E168d2/d2* and *-/-* retinas. The activity of these sites in *-/-* retinas suggests that they are re-used by another TF in cellular contexts where CRX levels are reduced or absent, and that this other TF is able to activate at the same binding sites where CRX represses (White et al. 2016). A probable candidate TF is the CRX ortholog OTX2, which plays a major role in retinal development and whose binding specificity is nearly identical to that of CRX (Furukawa et al. 2002; Nishida et al. 2003; Koike et al. 2007). These CREs thus may be CRX-bound silencers in rod photoreceptors (where CRX expression is high and OTX2 expression is low), and OTX2-bound enhancers in another cell type with high OTX2 and low CRX expression, such as bipolar cells (Koike et al. 2007; Murphy et al. 2019). CRX and OTX2, despite close homology in both their DNA binding and effector domains, have non-overlapping functions (Yamamoto et al. 2020; Terrell et al. 2012), which would allow them to play different regulatory roles at the same CREs. Currently there are a limited number of examples of CREs that act as silencers in one cell type and enhancers in another (Gisselbrecht et al. 2020). However, such bifunctional elements could be part of an effective regulatory strategy during the development of a population of multipotent progenitor cells into closely related cell types with overlapping regulators, as occurs in the retina (Swaroop et al. 2010; Murphy et al. 2019).

CREs bound by CRX differ in their chromatin environments, notably the cell-type specificity and developmental timing of chromatin accessibility (Ruzycki et al. 2018). We found that the activities of these CREs measured episomally by MPRA at P8 showed some correspondence with the chromatin environment at their native genomic loci. Distal CREs whose chromatin accessibility does not peak until 8 weeks after birth were depleted of strong enhancers and enriched for silencers, while proximal CREs that are already accessible by P0 contained the highest fraction of strong enhancers of any chromatin group. The enrichment of silencers among late-acting CRX-bound CREs suggests that these elements may have sequence-encoded repressive features that prevent premature activation. Our results indicate that episomal MPRAs capture some sequence-encoded features that determine differences in chromatin environment in the genome.

Many inherited human diseases are linked with variants in TFs (Tremblay et al. 2018; Herman et al. 2021; Sun and Chen 2023). To understand how different variants in the same TF gene produce different disease phenotypes, it is critical to understand the distinct effects of these variants on *cis*-regulatory function across a broad range of target CREs. Such regulome-wide effects of TF variants cannot be easily predicted from the biochemical details of interactions between TFs and co-factors, since these often vary in complex ways between CREs. Our work shows that MPRAs are an effective, direct readout of variant effects on TF function, and that they can yield insights into mechanisms of TF variants at differen target CREs. The approach described here thus complements the information learned from transcriptional and chromatin profiling assays.

## Methods

### Library Design and Cloning

The MPRA library contained 1,723 genomic sequences centered on CRX ChIP-seq peaks that overlapped ATAC-seq peaks from P14 retina (Ruzycki et al. 2018; Corbo et al. 2010). Sequences were 164 bp in length and represented in the library by four unique barcodes. As negative controls we included 100 scrambled sequences. Basal promoters were represented with 16 unique barcodes to ensure precise measurement of basal levels.

ChIP-seq peaks were sampled based on two sets of annotations described by Ruzycki et al. (Ruzycki et al. 2018) (Table 1). These annotations include 1) the presence or absence of histone marks H3K4me3 and H3K27ac in P14 wild-type retina, and 2) the loss or retention of chromatin accessibility in *Crx^-/-^* retina (*i.e.*, whether accessibility is CRX dependent or independent). For each of the wild-type ChIP peaks in the library, we created a mutant version in which all copies of the CRX motif with a p-value of 2.5 × 10^−3^ or less were mutated by changing the core TAAT to TACT (White et al. 2013; Friedman et al. 2021). To test the hypothesis that CRX mutants might alter cooperative activity at CREs, we selected 100 library sequences with only two copies of the CRX motif, and created single and double mutants.

The MPRA library was cloned following our previously described protocol (White et al. 2013; White et al. 2016; Friedman et al. 2021). Library sequences were synthesized by Agilent as barcoded oligonucleotides with the following structure: 5’ priming sequence (GTAGCGTCTGTCCGT)/EcoRI site/164 bp Library sequence/SpeI site/C/SphI site/Barcode sequence/NotI site/3’ priming sequence (CAACTACTACTACAG). The oligo library was amplified by four cycles of PCR using New England Biolabs (NEB) Phusion High-fidelity Master Mix. Amplicons were purified on a 2% agarose gel, digested with EcoRI-HF and NotI-HF (NEB), then cloned into the EagI and EcoRI sites of the plasmid pJK03 with T4 ligase (NEB). The libraries were transformed into 5-alpha electrocompetent cells (NEB). The plasmid library was prepared by maxi-prep (Sigma), digested with SphI-HF and SpeI-HF (NEB) and then treated with Antarctic phosphatase (NEB). The basal promoter with *DsRed* was amplified from pJK01 (*RHO*) (Kwasnieski et al. 2012) and purified from a 1% agarose gel. The basal promoter amplicon was digested with NheI-HF and SphI-HF, then cloned into the libraries with T4 ligase (NEB). The library was transformed into 5-alpha electrocompetent cells (NEB), grown in liquid culture, then prepared by maxi-prep (Sigma).

### Retinal Explant Electroporation and Library Sequencing

All mouse lines were on the C57BL/6J background and free of rd1 and rd8 mutations. The Crx^R90W/W^, Crx^R90W/+^, Crx^E168d2/d2^and Crx^E168d2/+^ lines and their genotyping procedures were described by Tran et al. (Tran et al. 2014). All procedures involving mice were approved by the Animal Studies Committee of Washington University in St. Louis and performed under Animal Welfare Assurance # A-3381-01 and Protocols # 21-0414 (to SC). Experiments were carried out in strict accordance with recommendations in the Guide for the Care and Use of Laboratory Animals of the National Institutes of Health (Bethesda, MD), the Washington University Policy on the Use of Animals in Research, and the Guidelines for the Use of Animals in Visual Research of the Association for Research in Ophthalmology and Visual Sciences. Every effort was made to minimize the animals’ suffering, anxiety and discomfort.

Retinal explant electroporations were conducted as previously described (Kwasnieski et al. 2012). Retinas were isolated from P0 mice and electroporated with 30 µg MPRA library and 30 µg pCAG-GFP as a transfection control. After culturing for 8 days retinas were placed in TRIzol (ThermoFisher) and RNA was extracted. Three replicate electroporations (3 retinas each) were performed.

cDNA was prepared using Superscript Reverse Transcriptase III (ThermoFisher) with oligo dT primers following the manufacturer’s instructions. Barcodes were amplified from cDNA or input plasmid DNA using 24 cycles of PCR with Phusion High Fidelity Master Mix (NEB). Amplicons were purified with the Monarch PCR kit (NEB), digested with MfeI-HF and SphI-HF (NEB) and ligated to custom sequencing adapters with PE2 indexing barcodes and phased P1 barcodes. Following 20 cycles of enrichment PCR, libraries were sequenced on the Illumina NextSeq platform to greater than 1000x coverage per sample.

### MPRA Quantification and Analysis

All sequencing reads were processed regardless of quality score. Sequenced barcodes were matched to expected library members, and only reads with a sequenced barcode that perfectly matched an expected library barcode were counted. Barcode counts were normalized by reads per million (RPM) for each sample. To account for differences in barcode representation in the pooled library, RNA reads were normalized to input DNA reads (Kwasnieski et al. 2012). RNA/DNA ratios were averaged over all barcodes for each element, and across biological replicates. CREs were classified into one of five activity groups based on their activity relative to the basal promoter alone. Expression levels were approximately log-normally distributed, so we computed the log-normal parameters for each sequence and then performed Welch’s t-test. We corrected for multiple hypotheses using the Benjamini-Hochberg FDR procedure. We corrected for multiple hypotheses in each library separately to account for any potential batch effects between libraries. The log2 expression was calculated after adding a pseudocount of 1 × 10^−3^ to every sequence. CREs with a transcriptional activity that was not significantly different from basal were deemed “inactive”. Remaining CREs with an activity significantly greater than basal were deemed “strong enhancers” if their activity was greater than the 95th percentile of the 100 scrambled control sequences, and “weak enhancers” otherwise. “Strong silencers” and “weak silencers” were classified by having activity significantly lower than the basal promoter, with strong silencers less than the 5th percentile of the 100 scrambled controls.

The information content and predicted CRX occupancy of each CRE sequence was computed as previously described (Friedman et al. 2021). Briefly, CRX predicted occupancy is determined by using the CRX energy weight matrix to calculate the probability of CRX binding at every 8-mer subsequence of each CRE. The binding probabilities at each position are summed to give the predicted CRX occupancy for the entire 164 bp sequence. Predicted occupancies for eight lineage-specific TFs described in the main text were used to calculate information content scores as described above and in Friedman et al. 2021.

To analyze motif enrichment across CRE subgroups, SEA (“Simple Enrichment Analysis”, MEME Suite v5.5.2) was used with default settings and the HOCOMOCO Mouse v11 CORE motif database (Bailey and Grant 2021; Kulakovskiy et al. 2018). Analysis scripts and Python notebooks for the described analyses are available at https://github.com/barakcohenlab/retinopathy-manuscript, including the source code necessary to generate all data figure panels. The completed MPRA analysis with activity and classifications of each CRE can be found in Supplemental Table S1.

## Data Access

All raw and processed sequencing data generated in this study have been deposited at the NCBI Gene Expression Omnibus (GEO; https://www.ncbi.nlm.nih.gov/geo/) under accession number GSE230090.

## Acknowledgements

This work was supported by NIH grants R01 EY027784 (to BC and SC), R01 EY012543 (to SC), R01 GM121755 (to MAW), F30 EY033640 (to JLS), F31 HG011431 (to RZF), and Research to Prevent Blindness (to SC and DOVS).

## References

Bailey TL, Grant CE. 2021. SEA: Simple Enrichment Analysis of motifs. bioRxiv.

Berger W, Kloeckener-Gruissem B, Neidhardt J. 2010. The molecular basis of human retinal and vitreoretinal diseases. Progress in Retinal and Eye Research 29: 335–375.

Chau K-Y, Chen S, Zack DJ, Ono SJ. 2000. Functional Domains of the Cone-Rod Homeobox (CRX) Transcription Factor. Journal of Biological Chemistry 275: 37264–37270.

Chen S, Wang Q-L, Nie Z, Sun H, Lennon G, Copeland NG, Gilbert DJ, Jenkins NA, Zack DJ. 1997. Crx, a Novel Otx-like Paired-Homeodomain Protein, Binds to and Transactivates Photoreceptor Cell-Specific Genes. Neuron 19: 1017–1030.

Chen S, Wang Q-L, Xu S, Liu I, Li LY, Wang Y, Zack DJ. 2002. Functional analysis of conerod homeobox (CRX) mutations associated with retinal dystrophy. Human Molecular Genetics 11: 873–884.

Corbo JC, Lawrence KA, Karlstetter M, Myers CA, Abdelaziz M, Dirkes W, Weigelt K, Seifert M, Benes V, Fritsche LG, et al. 2010. CRX ChIP-seq reveals the cis-regulatory architecture of mouse photoreceptors. Genome Research 20: 1512–1525.

Creyghton MP, Cheng AW, Welstead GG, Kooistra T, Carey BW, Steine EJ, Hanna J, Lodato MA, Frampton GM, Sharp PA, et al. 2010. Histone H3K27ac separates active from poised enhancers and predicts developmental state. Proceedings of the National Academy of Sciences of the United States of America 107: 21931–21936.

DelRosso N, Tycko J, Suzuki P, Andrews C, Aradhana null, Mukund A, Liongson I, Ludwig C, Spees K, Fordyce P, et al. 2023. Large-scale mapping and mutagenesis of human transcriptional effector domains. Nature 616: 365–372.

Evans K, Fryer A, Inglehearn C, Duvall-Young J, Whittaker JL, Gregory CY, Butler R, Ebenezer N, Hunt DM, Bhattacharya S. 1994. Genetic linkage of cone-rod retinal dystrophy to chromosome 19q and evidence for segregation distortion. Nature Genetics 6: 210–213.

Freund CL, Gregory-Evans CY, Furukawa T, Papaioannou M, Looser J, Ploder L, Bellingham J, Ng D, Herbrick J-AS, Duncan A, et al. 1997. Cone-Rod Dystrophy Due to Mutations in a Novel Photoreceptor-Specific Homeobox Gene (CRX) Essential for Maintenance of the Photoreceptor. Cell 91: 543–553.

Freund CL, Wang Q-L, Chen S, Muskat BL, Wiles CD, Sheffield VC, Jacobson SG, Mclnnes RR, Zack DJ, Stone EM. 1998. De novo mutations in the CRX homeobox gene associated with Leber congenital amaurosis. Nature Genetics 18: 311–312.

Friedman RZ, Granas DM, Myers CA, Corbo JC, Cohen BA, White MA. 2021. Information content differentiates enhancers from silencers in mouse photoreceptors. eLife 10: e67403.

Fujinami-Yokokawa Y, Fujinami K, Kuniyoshi K, Hayashi T, Ueno S, Mizota A, Shinoda K, Arno G, Pontikos N, Yang L, et al. 2020. Clinical and Genetic Characteristics of 18 Patients from 13 Japanese Families with CRX-associated retinal disorder: Identification of Genotype-phenotype Association. Scientific Reports 10: 9531.

Furukawa A, Koike C, Lippincott P, Cepko CL, Furukawa T. 2002. The mouse Crx 5’-upstream transgene sequence directs cell-specific and developmentally regulated expression in retinal photoreceptor cells. The Journal of Neuroscience: The Official Journal of the Society for Neuroscience 22: 1640–1647.

Furukawa T, Morrow EM, Cepko CL. 1997. Crx, a Novel otx-like Homeobox Gene, Shows Photoreceptor-Specific Expression and Regulates Photoreceptor Differentiation. Cell 91: 531–541.

Gisselbrecht SS, Palagi A, Kurland JV, Rogers JM, Ozadam H, Zhan Y, Dekker J, Bulyk ML. 2020. Transcriptional Silencers in Drosophila Serve a Dual Role as Transcriptional Enhancers in Alternate Cellular Contexts. Molecular Cell 77: 324–337.e8.

Grant CE, Bailey TL, Noble WS. 2011. FIMO: Scanning for occurrences of a given motif. Bioinformatics 27: 1017–1018.

Hennig AK, Peng G-H, Chen S. 2008. Regulation of photoreceptor gene expression by Crx-associated transcription factor network. Brain Research 1192: 114–133.

Herman L, Todeschini A-L, Veitia RA. 2021. Forkhead Transcription Factors in Health and Disease. Trends in genetics: TIG 37: 460–475.

Hlawatsch J, Karlstetter M, Aslanidis A, Lückoff A, Walczak Y, Plank M, Böck J, Langmann T. 2013. Sterile alpha motif containing 7 (Samd7) is a novel crx-regulated transcriptional repressor in the retina. PloS One 8: e60633.

Huang L, Xiao X, Li S, Jia X, Wang P, Guo X, Zhang Q. 2012. CRX variants in conerod dystrophy and mutation overview. Biochemical and Biophysical Research Communications 426: 498–503.

Hughes AEO, Myers CA, Corbo JC. 2018. A massively parallel reporter assay reveals context-dependent activity of homeodomain binding sites in vivo. Genome Research 28: 1520– 1531.

Hull S, Arno G, Plagnol V, Chamney S, Russell-Eggitt I, Thompson D, Ramsden SC, Black GCM, Robson AG, Holder GE, et al. 2014. The phenotypic variability of retinal dystrophies associated with mutations in CRX, with report of a novel macular dystrophy phenotype. Investigative Ophthalmology & Visual Science 55: 6934–6944.

Jacobson SG, Cideciyan AV, Huang Y, Hanna DB, Freund CL, Affatigato LM, Carr RE, Zack DJ, Stone EM, McInnes RR. 1998. Retinal degenerations with truncation mutations in the cone-rod homeobox (CRX) gene. Investigative Ophthalmology & Visual Science 39: 2417– 2426.

Jeon CJ, Strettoi E, Masland RH. 1998. The major cell populations of the mouse retina. The Journal of Neuroscience: The Official Journal of the Society for Neuroscience 18: 8936– 8946.

Karousis ED, Mühlemann O. 2019. Nonsense-Mediated mRNA Decay Begins Where Translation Ends. Cold Spring Harbor Perspectives in Biology 11: a032862.

Kim YJ, Lee M, Lee Y-T, Jing J, Sanders JT, Botten GA, He L, Lyu J, Zhang Y, Mettlen M, et al. 2023. Light-activated macromolecular phase separation modulates transcription by reconfiguring chromatin interactions. Science Advances 9: eadg1123.

Koike C, Nishida A, Ueno S, Saito H, Sanuki R, Sato S, Furukawa A, Aizawa S, Matsuo I, Suzuki N, et al. 2007. Functional Roles of Otx2 Transcription Factor in Postnatal Mouse Retinal Development. Molecular and Cellular Biology 27: 8318–8329.

Koyanagi Y, Akiyama M, Nishiguchi KM, Momozawa Y, Kamatani Y, Takata S, Inai C, Iwasaki Y, Kumano M, Murakami Y, et al. 2019. Genetic characteristics of retinitis pigmentosa in 1204 Japanese patients. Journal of Medical Genetics 56: 662–670.

Kulakovskiy IV, Vorontsov IE, Yevshin IS, Sharipov RN, Fedorova AD, Rumynskiy EI, Medvedeva YA, Magana-Mora A, Bajic VB, Papatsenko DA, et al. 2018. HOCOMOCO: Towards a complete collection of transcription factor binding models for human and mouse via large-scale ChIP-Seq analysis. Nucleic Acids Research 46: D252–D259.

Kwasnieski JC, Mogno I, Myers CA, Corbo JC, Cohen BA. 2012. Complex effects of nucleotide variants in a mammalian cis-regulatory element. Proceedings of the National Academy of Sciences 109: 19498–19503.

Langouët M, Jolicoeur C, Javed A, Mattar P, Gearhart MD, Daiger SP, Bertelsen M, Tranebjærg L, Rendtorff ND, Grønskov K, et al. 2022. Mutations in BCOR , a co-repressor of CRX/OTX2 , are associated with early-onset retinal degeneration. Science Advances 8: eabh2868.

Lee J, Myers CA, Williams N, Abdelaziz M, Corbo JC. 2010. Quantitative fine-tuning of photoreceptor cis-regulatory elements through affinity modulation of transcription factor binding sites. Gene Therapy 17: 1390–1399.

Loell KJ, Friedman RZ, Myers CA, Corbo JC, Cohen BA, White MA. 2023. Transcription factor interactions explain the context-dependent activity of CRX binding sites. bioRxiv.

Mitton KP, Swain PK, Chen S, Xu S, Zack DJ, Swaroop A. 2000. The Leucine Zipper of NRL Interacts with the CRX Homeodomain. Journal of Biological Chemistry 275: 29794–29799.

Murphy DP, Hughes AE, Lawrence KA, Myers CA, Corbo JC. 2019. Cis-regulatory basis of sister cell type divergence in the vertebrate retina. eLife 8: e48216.

Ng CC, Carrera WM, McDonald HR, Agarwal A. 2020. Heterozygous CRX R90W mutation-associated adult-onset macular dystrophy with phenotype analogous to benign concentric annular macular dystrophy. Ophthalmic Genetics 41: 485–490.

Nichols LL, Alur RP, Boobalan E, Sergeev YV, Caruso RC, Stone EM, Swaroop A, Johnson MA, Brooks BP. 2010. Two novel CRX mutant proteins causing autosomal dominant Leber congenital amaurosis interact differently with NRL. Human Mutation 31.

Nishida A, Furukawa A, Koike C, Tano Y, Aizawa S, Matsuo I, Furukawa T. 2003. Otx2 homeobox gene controls retinal photoreceptor cell fate and pineal gland development. Nature Neuroscience 6: 1255–1263.

Nishiguchi KM, Kunikata H, Fujita K, Hashimoto K, Koyanagi Y, Akiyama M, Ikeda Y, Momozawa Y, Sonoda K-H, Murakami A, et al. 2020. Association of CRX genotypes and retinal phenotypes confounded by variable expressivity and electronegative electroretinogram. Clinical & Experimental Ophthalmology 48: 644–657.

Omori Y, Kubo S, Kon T, Furuhashi M, Narita H, Kominami T, Ueno A, Tsutsumi R, Chaya T, Yamamoto H, et al. 2017. Samd7 is a cell type-specific PRC1 component essential for establishing retinal rod photoreceptor identity. Proceedings of the National Academy of Sciences of the United States of America 114: E8264–E8273.

Peng G-H, Ahmad O, Ahmad F, Liu J, Chen S. 2005. The photoreceptor-specific nuclear receptor Nr2e3 interacts with Crx and exerts opposing effects on the transcription of rod versus cone genes. Human Molecular Genetics 14: 747–764.

Peng G-H, Chen S. 2007. Crx activates opsin transcription by recruiting HAT-containing co-activators and promoting histone acetylation. Human Molecular Genetics 16: 2433–2452.

Perrault I, Hanein S, Gerber S, Barbet F, Dufier J-L, Munnich A, Rozet J-M, Kaplan J. 2003. Evidence of autosomal dominant Leber congenital amaurosis (LCA) underlain by a CRX heterozygous null allele. Journal of Medical Genetics 40: e90.

Rister J, Razzaq A, Boodram P, Desai N, Tsanis C, Chen H, Jukam D, Desplan C. 2015. Singlebase pair differences in a shared motif determine differential Rhodopsin expression. Science 350: 1258–1261.

Rivolta C, Berson EL, Dryja TP. 2001. Dominant Leber congenital amaurosis, cone-rod degeneration, and retinitis pigmentosa caused by mutant versions of the transcription factor CRX. Human Mutation 18: 488–498.

Roduit R, , Schorderet DF. 2009. Mutations in the DNA-Binding Domain of NR2E3 Affect In Vivo Dimerization and Interaction with CRX ed. O.J. Manzoni. PLoS ONE 4: e7379.

Ruzycki PA, Tran NM, Kolesnikov AV, Kefalov VJ, Chen S. 2015. Graded gene expression changes determine phenotype severity in mouse models of CRX-associated retinopathies. Genome Biology 16.

Ruzycki PA, Zhang X, Chen S. 2018. CRX directs photoreceptor differentiation by accelerating chromatin remodeling at specific target sites. Epigenetics & Chromatin 11.

Sohocki MM, Sullivan LS, Mintz-Hittner HA, Birch D, Heckenlively JR, Freund CL, McInnes RR, Daiger SP. 1998. A Range of Clinical Phenotypes Associated with Mutations in CRX, a Photoreceptor Transcription-Factor Gene. The American Journal of Human Genetics 63: 1307–1315.

Srinivas M, Ng L, Liu H, Jia L, Forrest D. 2006. Activation of the blue opsin gene in cone photoreceptor development by retinoid-related orphan receptor beta. Molecular Endocrinology (Baltimore, Md) 20: 1728–1741.

Sun C, Chen S. 2023. Disease-causing mutations in genes encoding transcription factors critical for photoreceptor development. Frontiers in Molecular Neuroscience 16: 1134839.

Swain PK, Chen S, Wang QL, Affatigato LM, Coats CL, Brady KD, Fishman GA, Jacobson SG, Swaroop A, Stone E, et al. 1997. Mutations in the cone-rod homeobox gene are associated with the cone-rod dystrophy photoreceptor degeneration. Neuron 19: 1329–1336.

Swaroop A, Kim D, Forrest D. 2010. Transcriptional regulation of photoreceptor development and homeostasis in the mammalian retina. Nature Reviews Neuroscience 11: 563–576.

Swaroop A, Wang Q-L, Wu W, Cook J, Coats C, Xu S, Chen S, Zack DJ, Sieving PA. 1999. Leber Congenital Amaurosis Caused by a Homozygous Mutation (R90W) in the Homeodomain of the Retinal Transcription Factor CRX: Direct Evidence for the Involvement of CRX in the Development of Photoreceptor Function. Human Molecular Genetics 8: 299– 305.

Terrell D, Xie B, Workman M, Mahato S, Zelhof A, Gebelein B, Cook T. 2012. OTX2 and CRX rescue overlapping and photoreceptor-specific functions in the Drosophila eye. Developmental Dynamics: An Official Publication of the American Association of Anatomists 241: 215–228.

Tran NM, Chen S. 2014. Mechanisms of blindness: Animal models provide insight into distinct CRX-associated retinopathies. Developmental Dynamics 243: 1153–1166.

Tran NM, Zhang A, Zhang X, Huecker JB, Hennig AK, Chen S. 2014. Mechanistically Distinct Mouse Models for CRX-Associated Retinopathy ed. G.S. Barsh. PLoS Genetics 10: e1004111.

Tremblay M, Sanchez-Ferras O, Bouchard M. 2018. GATA transcription factors in development and disease. Development (Cambridge, England) 145: dev164384.

Vermeulen M, Eberl HC, Matarese F, Marks H, Denissov S, Butter F, Lee KK, Olsen JV, Hyman AA, Stunnenberg HG, et al. 2010. Quantitative interaction proteomics and genome-wide profiling of epigenetic histone marks and their readers. Cell 142: 967–980.

White MA, Kwasnieski JC, Myers CA, Shen SQ, Corbo JC, Cohen BA. 2016. A Simple Grammar Defines Activating and Repressing cis-Regulatory Elements in Photoreceptors. Cell Reports 17: 1247–1254.

White MA, Myers CA, Corbo JC, Cohen BA. 2013. Massively parallel in vivo enhancer assay reveals that highly local features determine the cis-regulatory function of ChIP-seq peaks. Proceedings of the National Academy of Sciences 110: 11952–11957.

Wilken MS, Brzezinski JA, La Torre A, Siebenthall K, Thurman R, Sabo P, Sandstrom RS, Vierstra J, Canfield TK, Hansen RS, et al. 2015. DNase I hypersensitivity analysis of the mouse brain and retina identifies region-specific regulatory elements. Epigenetics & Chromatin 8: 8.

Wilson D, Sheng G, Lecuit T, Dostatni N, Desplan C. 1993. Cooperative dimerization of paired class homeo domains on DNA. Genes & Development 7: 2120–2134.

Yamamoto H, Kon T, Omori Y, Furukawa T. 2020. Functional and Evolutionary Diversification of Otx2 and Crx in Vertebrate Retinal Photoreceptor and Bipolar Cell Development. Cell Reports 30: 658–671.e5.

Yanagi Y, Masuhiro Y, Mori M, Yanagisawa J, Kato S. 2000. P300/CBP acts as a coactivator of the cone-rod homeobox transcription factor. Biochemical and Biophysical Research Communications 269: 410–414.

Zagozewski JL, Zhang Q, Pinto VI, Wigle JT, Eisenstat DD. 2014. The role of homeobox genes in retinal development and disease. Developmental Biology 393: 195–208.

Zhao S, Hong CKY, Myers CA, Granas DM, White MA, Corbo JC, Cohen BA. 2023. A single-cell massively parallel reporter assay detects cell-type-specific gene regulation. Nature Genetics 55: 346–354.

Zheng Y, Sun C, Zhang X, Ruzycki P, Chen S. 2023. Missense mutations in CRX homeodomain cause dominant retinopathies through two distinct mechanisms. bioRxiv: The Preprint Server for Biology 2023.02.01.526652.

Ziviello C, Simonelli F, Testa F, Anastasi M, Marzoli SB, Falsini B, Ghiglione D, Macaluso C, Manitto MP, Garrè C, et al. 2005. Molecular genetics of autosomal dominant retinitis pigmentosa (ADRP): A comprehensive study of 43 Italian families. Journal of Medical Genetics 42: e47.

