## Supplementary figures and images for "Pathogenic variants in Crx have distinct cis-regulatory effects on enhancers and silencers in photoreceptors"

### Supplemental Figure 1

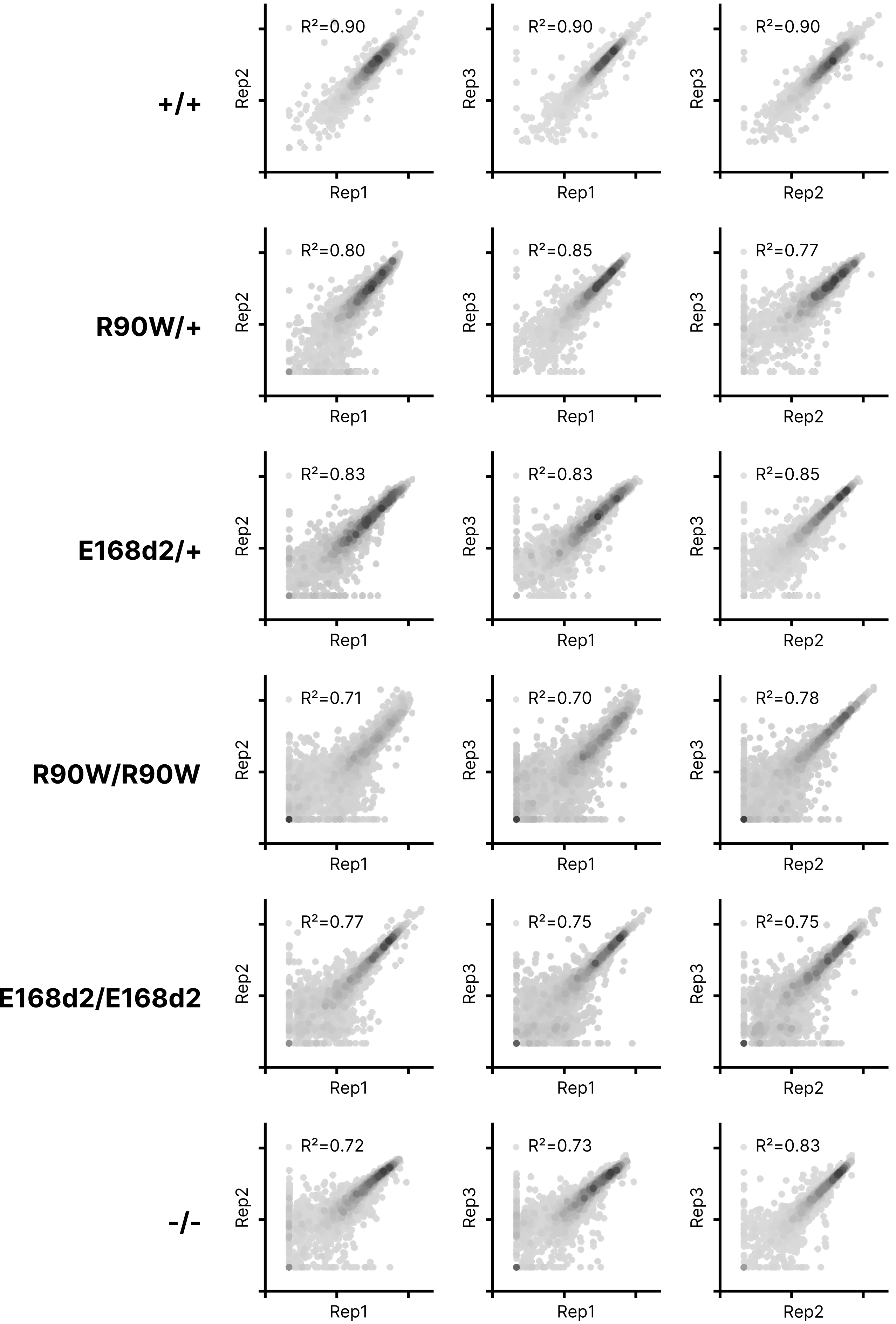

### Supplemental Figure 2

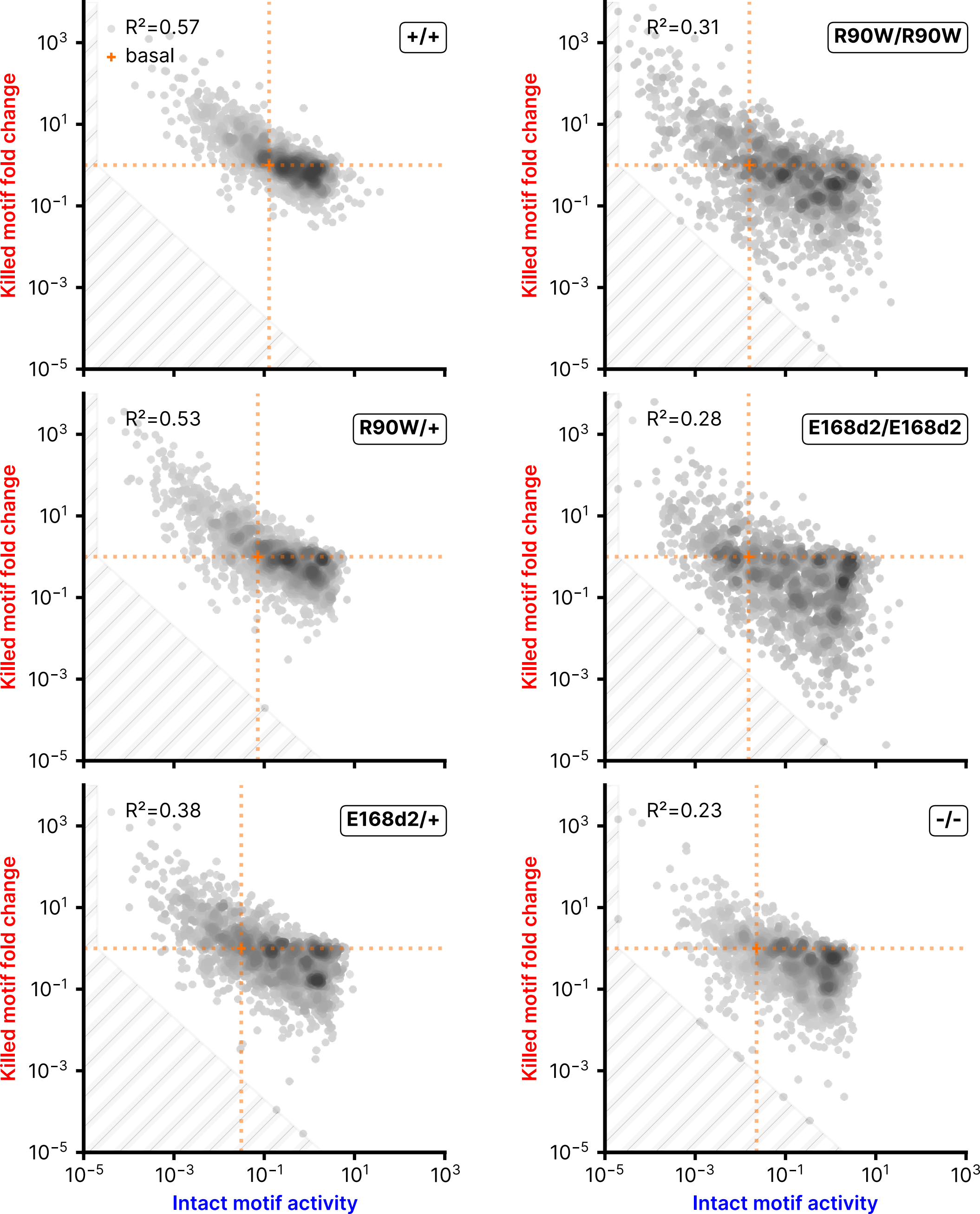

### Supplemental Figure 3

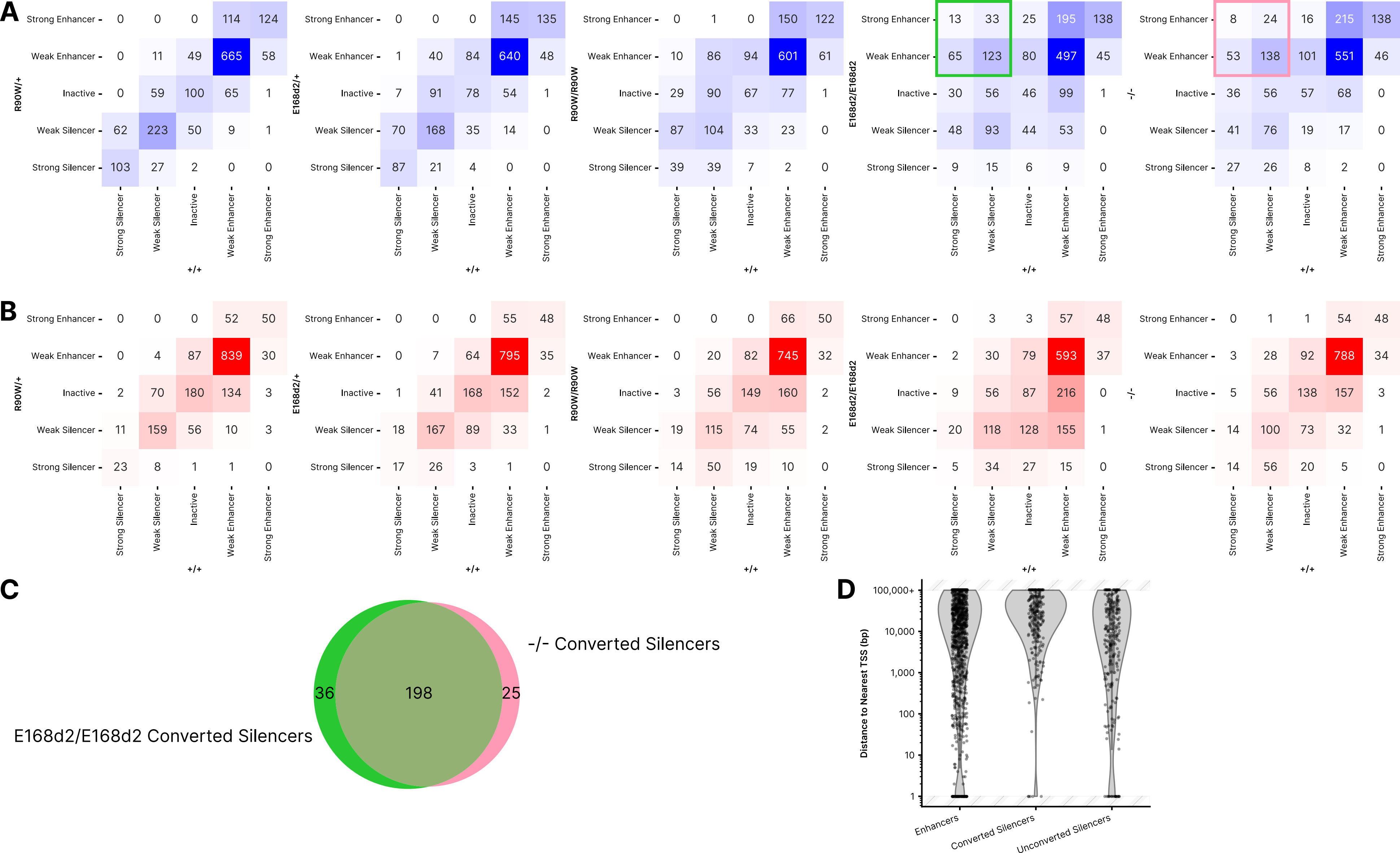

### Supplemental Figure 4

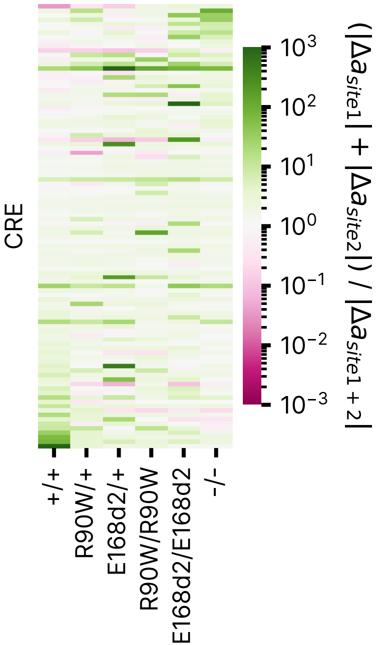
